# Temozolomide Induces Aberrant RNA Alkylation and Widespread Translational Repression

**DOI:** 10.64898/2026.09.01.748722

**Authors:** Jayden Rhodes, Abigail Grace Johnston, Katherine Marlow Brazell, Xisheng Liu, John L. Hartman, Zhangli Su

## Abstract

Temozolomide (TMZ) is a frontline alkylating chemotherapy, yet its direct impact on RNA modification and global translation dynamics remains poorly understood. Here, we demonstrate that TMZ induces pervasive RNA alkylation causing severe translational impairment. TMZ directly deposits aberrant methyl groups onto single-stranded mRNA *in vitro*, creating physical lesions that lower translational efficiency. In glioblastoma cells, acute TMZ exposure triggers a rapid, widespread accumulation of m^7^G on cellular RNAs, leading to the significant attenuation of global protein synthesis. Nanopore direct RNA sequencing identified distinct guanine-specific error signatures and sequence context preferences associated with TMZ-induced damage. Using a quantitative yeast spike-in ribosome profiling strategy, we mapped this translational repression at transcript-level, revealing a global downregulation of translational efficiency. This widespread repression disproportionately targets highly interconnected networks essential for cellular proliferation, specifically chromosome organization. We show that the severity of this translational repression is driven by a transcript’s coding guanine density, stability and translation initiation speed. Together, our findings suggest that TMZ-induced alkylation targets stable, highly translated, guanine-rich transcripts. This establishes aberrant RNA methylation and subsequent translational arrest as a potential mechanism of temozolomide cytotoxicity.

## Introduction

Glioblastoma (GBM) is a fast growing and aggressive malignancy of the central nervous system. Despite a rigorous standard of care including surgery, radiation, and chemotherapy, patient prognosis remains extremely poor, with median survival rarely exceeding 15 months post-diagnosis (1–5). This high mortality is largely driven by tumor recurrence and treatment resistance, including to the primary therapeutic agent, temozolomide (TMZ).

Temozolomide (TMZ) is an oral alkylating agent capable of crossing the blood brain barrier (6,7). Patients take up to 200 mg/m^2^ TMZ in capsule form daily during maintenance therapy (8). Under physiological conditions, TMZ spontaneously hydrolyzes and decomposes to form a methyldiazonium ion, a highly reactive intermediate that deposits methyl groups onto nucleotide bases (7,9). Historically, the cytotoxic effect of TMZ has been attributed almost entirely to its ability to induce aberrant DNA methylation, specifically at the *N*7 and *O*6 positions of guanine and the *N*3 position of adenine (6,10). Methylation at these sites triggers DNA replication collapse, ultimately leading to apoptosis (7). In *in vitro* settings, researchers frequently expose GBM cell lines to high concentrations of TMZ, often up to 1 mM, to model severe chemoresistance or map acute alkylation damage (11,12). Specifically, it was found that TMZ prefers certain trinucleotide context for *O*6-meG DNA alkylation (11). Because TMZ hydrolyzes rapidly, direct DNA alkylation occurs within the first few hours of exposure; however, downstream markers of catastrophic DNA damage, such as double-strand breaks and γH2AX foci, typically require 24 to 72 hours to manifest as cells attempt to replicate (13). On the other hand, RNA is significantly more abundant than DNA in the cell and is predominantly single-stranded, lacking the protection of a complementary strand or stable histone packaging. This structural accessibility renders cellular RNA highly susceptible to alkylation damage, yet the impact of TMZ on RNA substrates has been largely overshadowed by DNA-centric models.

Alkylating agents are prevalent as both environmental contaminants and frontline clinical treatments. Emerging evidence highlights that exogenous alkylating agents can introduce aberrant RNA modifications and severely disrupt RNA folding, stability, and translational fidelity (14–16). For example, the alkylating chemical methyl methanesulfonate (MMS) preferentially methylate RNA more than DNA (17), creating modifications like m^1^A (*N*1-methyladenosine), m^3^C (*N*3-methylcytidine) and m^7^G (*N*7-methylguanosine) (18). These damage-induced modifications often disrupt Watson-Crick base pairing or introduce positive charges. Furthermore, recent findings demonstrate that this rapid burst of RNA alkylation serves as an early cellular sensor to trigger the downstream DNA damage response (14). In contrast to these nitrogen-based modifications, alkylation at the oxygen position of guanine (*O*6-methylguanosine) is the hallmark cytotoxic lesion induced by TMZ on DNA (19). Building on the inherent vulnerability of RNA, we reason that TMZ-mediated RNA alkylation could occur at high frequencies, contributing to its overall cellular impact. Interestingly, these damage-induced modifications are chemically identical to the endogenously catalyzed modifications. For instance, m^1^A, m^3^C and m^7^G are enriched on tRNAs to support tRNA folding, with m^1^A and m^7^G on specific internal mRNA sites with low stoichiometry (20,21). This is distinct from m^6^A (*N*6-methyladenosine), the most abundant internal mRNA modification, which preserves standard Watson-Crick base pairing to regulate transcript turnover (22).

In this study, we set out to determine the extent and functional consequences of aberrant RNA methylation induced by the chemotherapy reagent TMZ. We demonstrate that *in vitro* TMZ exposure results in pervasive RNA alkylations including m^1^A, m^3^C and m^7^G modifications, which significantly impair mRNA translation. Consistent with these biochemical findings, acute TMZ treatment in GBM cells attenuates global protein synthesis alongside a significant induction of global m^7^G RNA methylation. Nanopore direct RNA sequencing identified distinct guanine-specific error signatures and sequence context preferences (such as GGG and AGG motifs) associated with TMZ-induced damage both in vitro and in cellular transcripts. Using quantitative yeast spike-in translatome profiling, we reveal the specific features correlating with this widespread translational repression by TMZ treatment. The severity of this TMZ-induced translational repression is driven by a transcript’s coding high guanine density, basal stability and rapid translation initiation kinetics. Together, these results elucidate the direct impact of TMZ on the transcriptome, highlighting aberrant RNA modification and subsequent translational arrest as a critical, early driver of chemotherapeutic cytotoxicity.

## Materials and Methods

### In Vitro Treatment and RNA Repurification

1 µg of luciferase control RNA (Promega L4561) was treated with 30 mM final concentration of TMZ or MMS at room temperature for 30 minutes. *TMZ treatment:* 8.82 µL of 170 mM temozolomide (TMZ) (Sigma-Aldrich T2577-25MG) dissolved in DMSO was added to RNA in the presence of 0.5 µL of RNase inhibitor (Thermo Fisher Scientific AM2696), 2.5 µL of 1 M TE Buffer (Fisher Scientific 50-111-8116) in a total volume of 50 µL. As vehicle control, 8.82 µL of Dimethyl sulfoxide (DMSO) (Fisher Scientific BP231-100) was used. *MMS treatment:* 15 µL of 100 mM Methyl methanesulfonate (MMS) (Fisher Scientific 66-27-3) in water was added to RNA in the presence of 0.5 µL of RNase inhibitor (Thermo Fisher Scientific AM2696), 2.5 µL of 1 M TE Buffer (Fisher Scientific 50-111-8116) in a total volume of 50 µL. As vehicle control, 15 µL of water was used. After incubation, the RNA clean and concentrator kit (Zymo Research RCC-5) was used to purify the RNA for two rounds. To account for potential RNA loss during incubation and clean-up, a “No Treatment” condition was created by mixing 1 µg of luciferase control RNA with 49 µL of RNase-free H_2_O. This fifth condition was vortexed and remained on ice until all five of the samples’ RNA concentrations were read using a NanoDrop spectrophotometer. RNA integrity was checked by BioAnalyzer Pico Analysis at UAB Heflin Genomics Core.

### In Vitro Translation and Luciferase Assay

The Rabbit Reticulocyte Lysate System (Promega RL4960) was used to measure *in vitro* translation of the purified RNA species. 0.5 µL of RNase inhibitor, 9 µL of Rabbit Reticulocyte Lysate, 10 ng of one of the five different RNA treatment conditions, and RNase-free H_2_O up to a final volume 20 µL was mixed in a PCR tube and placed on ice. The samples were then heated in a PCR machine for 30 minutes at 30 °C and immediately transferred to −80 °C, where they remained overnight to quench the reaction. The next day, the samples were thawed placed in a 96-well plate (Fisher Scientific 07200589) before luminescence was read. The luminescence was quantified by the Promega GloMax luminometer with 10 seconds read time per well. LARII reagent (Promega PRE1980) was injected to each well before the reading.

### GBM Cell Culture, Treatment and RNA Extraction

U251-GM cells were purchased (Millipore Sigma 09063001-1VL) and maintained in HyClone Dulbecco’s High Glucose Modified Eagles medium with L-glutamine (Cyvita 16750-074) plus 10% fetal bovine serum (Cyvita SH3091003) and 1% penicillin/streptomycin (Gibco 15140122). Mycoplasma contamination was routinely checked by mycostrip kit (InvivoGen rep-mysnc-50). Cells were grown in humidified incubators with 5% CO2 at 37°C. 400,000 U251 cells were seeded in 6-cm dish and treated with 1 mM final concentration of temozolomide (TMZ) (Sigma-Aldrich T2577-25MG), Dimethyl sulfoxide (DMSO) (Fisher Scientific BP231-100), methyl methanesulfonate (MMS) diluted in water (Fisher Scientific 66-27-3) or mock treatment for 2 hours. After treatment, cells were subjected to RNA extraction or protein lysate collection. U251- GM cells were washed with cold 1X PBS two times before being resuspended in 500 µL Trizol reagent (Life Technologies 15596018). Total RNA was extracted by Directzol RNA Miniprep Plus kit with on-column DNase I treatment (ZYMO R2071) and eluted in 50 µL RNase-free water.

### Mass Spectrometry Analysis for Modified Nucleosides

RNA samples were submitted to UAB’s Targeted Metabolomics and Proteomics Laboratory (TMPL) for mass spectrometry, according to published protocols (23,24). S1 Nuclease (Thermo Fisher Scientific 18001016) was diluted to 10 units/µL by combining 1 µL of the Nuclease, 1 µL of 10X Reaction Buffer, and 8 µL of RNase-free H_2_O. 2.5 µg of the *in vitro* or cellular treated RNA species, 1 µL of diluted S1 Nuclease, 4 µL of 10X Reaction Buffer, and various amounts of RNase-free H_2_O was then added together to a total volume to 25 µL. The samples were mixed and incubated at 37 °C in a PCR machine overnight. The next day, 5 µL 10X FastAP Buffer (Thermo Fisher Scientific FEREF0651), 2 µL FastAP Enzyme (Thermo Fisher Scientific EF0651), and 3 µL of RNase-free H_2_O was mixed into the samples before being incubated at 37 °C for 1 hour. To remove the enzyme and retain the mononucleoside, the 50 µL samples were added to centrifugal devices with 3K Omega Filter Membranes (Cytiva 29300-606) and centrifuged at 10,000 x g for 20 minutes at 4 °C. The filtrates transferred to low DNA-binding tubes before submission to the TMPL core. Relative standard curves for each nucleoside were created: adenosine (Millipore Sigma A9251), cytidine (Millipore Sigma C122106), uridine (Millipore Sigma U3750), guanosine (Millipore Sigma G6752), *N*1- methyladenosine (Cayman Chemical 16937), *N*6-methyladenosine (Selleck Chemical S3190), *N*3-methylcytidine (Cayman Chemical 21064), *N7*-methylguanosine (Cayman Chemical 15988).

Methylated and unmethylated RNA nucleotides were quantified using electrospray ionization-liquid chromatography-tandem mass spectrometry (ESI-LC-MS/MS) on an SCIEX 7500+ Triple Quadrupole mass spectrometer (Sciex, Foster City, CA) coupled to a Exion AE UPLC system with a refrigerated autosampler AE (Sciex, Foster City, CA). Chromatographic separation was performed on an Atlantis T3, 3 µ column (80 Å, 100 × 2.1 mm; Waters, Milford, MA) maintained at 40 °C. The mobile phases consisted of water with 0.1% formic acid (A) and acetonitrile with 0.1% formic acid (B), delivered at 0.5 mL/min with a gradient from 2-15% B over 2.0 min, 15-98% to 3.0 min with a 0.5 min hold at 98%, followed by column flushing with 100% mobile phase B to 5 min. The injection volume was 5 µL, and the total run time was 5 min. Mass spectrometric detection was performed in positive electrospray ionization mode with an ion spray voltage of 5000 V, interface temperature of 500 °C, and gas settings of 40, 35, and 70 psi for curtain, GS1, and GS2, respectively. Multiple reaction monitoring (MRM) transitions were monitored as the following (EP – Entrance Potential, CE – Collision Energy, CXP – Collision Cell exit Potential):

| Compound | Q1<br>(Da) | Q2<br>(Da) | Dwell<br>(ms) | EP<br>(V) | CE<br>(V) | CXP<br>(V) |
| --- | --- | --- | --- | --- | --- | --- |
| *N*1-methyladenosine * | 282.1 | 150 | 10 | 10 | 25 | 10 |
| *N*6-methyladenosine ** | 282.1 | 150 | 10 | 10 | 25 | 10 |
| *N*3-methylcytidine | 258.1 | 126.2 | 10 | 10 | 20 | 10 |
| *N*7-methylguanosine *** | 298 | 166 | 10 | 10 | 15 | 10 |
| Guanosine | 284.1 | 152.1 | 10 | 10 | 15 | 10 |
| Adenosine | 268.1 | 136 | 10 | 10 | 25 | 10 |
| Cytidine | 244.1 | 112.1 | 10 | 10 | 15 | 10 |
| Uridine | 245.1 | 113 | 10 | 10 | 30 | 10 |
| * | Retention time at 1.03 min |  |  |  |  |  |
| ** | Retention time at 1.84 min |  |  |  |  |  |
| *** | Retention time at 1.07 min |  |  |  |  |  |

Data acquisition and analysis were performed using Analyst OS (version 3.4) and MultiQuant software (version 3.0) for respectively (SCIEX, Foster City, CA).

### m7G Dot Blot

35 ng of the *in vitro* treated luciferase RNAs and the control RNA were heated in a PCR machine for 2 minutes at 70 °C and immediately placed on ice. The samples were dried on the Hybond+ Nylon membrane (Avantor 95038-376) at room temperature for 2 minutes, before UV crosslink (1,200 uJ x 100). Blocking was completed using 3% milk in 1X PBS (Fisher Scientific BP3991) containing 0.05% Tween-20 (Fisher Scientific BP337-100) on an orbital shaker at room temperature for 30 minutes. The membrane was then probed with 1:1,000 primary m^7^G antibody (MBL RN017M, clone 4141-13) in 5% BSA/PBST on an orbital shaker overnight at 4 °C and 1:10,000 goat-anti-mouse secondary antibody (Cell Signaling #7076V) on an orbital shaker at room temperature for 1 hour. The membrane was incubated with HRP Enhanced Chemiluminescent Substrate (Sigma-Aldrich WBKLS0500) and imaged on BioRad Gel Documentation System.

### Puromycin Labeling and Western Blot

After TMZ and control treatment, puromycin labeling was performed to measure global protein translation by replacing media with 500 µM puromycin final concentration and incubated for 10 minutes before moving to collection. The cells were washed with 1X PBS and then lysed using 250 µL of RIPA Buffer with 100X Proteinase Inhibitor. The concentrations of the proteins were measured using the Qubit Protein Broad Range Assay (Thermo Fisher Scientific A50668).

10 µg protein lysates were separated by using a 4-20% Tris-Glycine gel (Thermo Fisher Scientific XP04205BOX) and transferred to Protran nitrocellulose membrane (Amersham 10063-173) via BioRad Semi-dry transfer. Blocking was performed using 3% milk in PBST. Primary antibody against puromycin (Millipore Sigma MABE343) was used at 1:2,000 in blocking buffer. For ATF-4 and HSP90, U251 cells were not labeled by puromycin labeling. 20 µg protein lysates were probed with 1:1,000 ATF4 antibody (Cell Signaling Technology 11815) or 1:2,000 HSP-90 antibody (Santa Cruz sc-13119). The Immobilon Western Chemiluminescent HRP system (Sigma-Aldrich WBKLS0500) was used for detection and imaged on BioRad Gel Documentation System.

### Nanopore Direct RNA Sequencing and Bioinformatics

The library was prepared from 100 ng firefly luciferase RNA or 1 µg of total U251 RNA according to the Oxford Nanopore technology (ONT) direct RNA sequencing protocol version DRS_9195_v4_revI_30Jul2025. This protocol specifically captures poly-A RNA species. The library was loaded onto the ONT FLO-MIN004RA flow cell and sequenced on a MinION Mk1B using MinKnow. Raw pod5 files were basecalled using Dorado v2.1.0 with the rna004_hac@v6.0.0 model. The resulting bam files were mapped using EPI2ME wf-alignment v1.2.6 using luciferase RNA sequence or gencode human transcriptome reference v50. We report mapped reads: (1) control firefly luciferase: 1395 reads, (2) *in vitro* 120 mM MMS treatment: 6889 reads, (3) *in vitro* 30 mM TMZ treatment: 6263 reads, (4) U251 DMSO treatment: 1,502,728 reads, (5) U251 1mM TMZ treatment: 965,784 reads.

After mapping, EpiNano was used to quantify basecalling error per position. For U251 data, the top 10 expressed protein-coding transcripts (ENST00001145575.1, ENST00000361739.1, ENST00000362079.2, ENST00000361624.2, ENST00000361227.2, ENST00000361899.2, ENST00000361453.3, ENST00000361381.2, ENST00000361390.2, and ENST00000368719.9) were selected for downstream analysis. (*1*) *<u>Delta error per position:</u>* Delta error was calculated by subtracting the control error rate from the TMZ or MMS treated error rate for each corresponding position, where the error rate per position was summed from mismatch, insertion, and deletion frequencies. Statistical significance for the resulting violin plots was determined using one-sample t-tests against a mean of zero for each base. We applied a minimum coverage cutoff of 50 reads for both control and treated samples across all analyzed positions. For the landscape plots, positions exhibiting a delta error greater than or equal to 5 percent were highlighted along the transcript length. A loess smoothed trendline was applied to the landscape data. (*2*) *<u>Motif analysis for in vitro TMZ treatment:</u>* We identified specific foreground and background Guanine (G) sites for motif analysis. Foreground G sites were defined as positions with a minimum coverage of 50 in both control and treated conditions with an error increase of at least 3 percent. Background G sites were defined as positions with a minimum coverage of 50 and an error increase of less than 1 percent. We extracted 3-mers using a window size of 1 for both foreground and background groups to identify unbiased motifs. This creates a 3-mer perfectly centered on the target site. Motif frequencies were compared using Fisher’s Exact Test. The resulting p-values were adjusted for multiple testing using the False Discovery Rate (FDR) method. Motifs were classified as significantly enriched or depleted based on an FDR of less than 0.05 and their log2 fold enrichment. (*3*) *<u>3-mer aggregate analysis</u> <u>for in vitro TMZ treatment:</u>* 3-mers centered on all G positions with coverage greater than 50 were extracted. The mean error increase between TMZ and control samples was calculated for motifs occurring at least 5 times. Z-scores were computed by comparing each motif mean error rate to the global mean error rate across all G sites. Significance was determined using FDR adjusted p-values. This classified motifs into distinct statistical tiers. (*4*) *<u>3-mer aggregate</u> <u>analysis for U251 TMZ treatment:</u>* Delta error between TMZ and DMSO treatments was calculated for the top 10 expressed protein-coding transcripts on positions with a minimum coverage of 50. A 3-base rolling average was applied to the delta error to generate a smoothed error metric. We extracted 3-mers using a window size of 1. The mean smoothed error was calculated for 3-mers occurring at least 20 times. Z-scores were computed by comparing the mean smoothed error of each 3-mer against the global mean smoothed error across all analyzed transcript positions. Significance was assigned using FDR adjusted p-values. The top 20 error prone 3-mers were selected for visualization.

### Ribo-seq and RNA-seq Library Preparation with Yeast Spike-in

The Ribo-seq was performed by combining Ribo-seq protocol with yeast spike-in normalization (25,26). (*1*) *<u>Mammalian and Yeast lysate preparation:</u>* Briefly, U251 cells were washed with ice-cold DPBS containing 100 µg/mL cycloheximide (CHX), scraped into 1x lysis buffer (20 mM Tris, pH 7.4, 150 mM NaCl, 5 mM MgCl_2_, 1mM DTT, 25 U/mL Turbo DNase I, 1% Triton X-100, 1x protease inhibitor cocktail), homogenized via a 26-gauge needle, and incubated on ice for 10 minutes. Lysates were clarified (15,000 × g, 10 min, 4°C). Flash-frozen yeast pellets were lysed via bead beating (0.5 mm glass beads, Biobasic) in 1x lysis buffer for 1 minute, clarified (20,000 × g, 5 min, 4°C), diluted to ∼7 A260 units/mL by Nanodrop, and stored at −80°C. Yeast lysate was added to mammalian lysates at 2% (w/w) based on Qubit Broad Range RNA measurements. For inputs, 10% of the combined lysate was saved for making RNA-seq libraries.

(*2*) *<u>RPF (ribosome protected fragments) preparation:</u>* Lysate RNA was quantified by Qubit RNA Broad Range kit (Invitrogen Q10211) and ∼10 µg aliquots were digested with 0.375 U/µg RNase I (Biosearch Technologies, N6901K) for 50 min at room temperature (20 - 25°C). Digestion was quenched with 200 U SUPERase•In (Invitrogen AM2696). Monosomes were isolated using pre-equilibrated MicroSpin S-400 HR columns (Cytiva GS27-5140-01) by centrifugation (600 × g, 2 minutes). RPFs were extracted from the eluates using acid phenol:chloroform (Invitrogen AM9720), followed by overnight precipitation at −20°C with 0.1 volumes of 3 M sodium acetate (pH 5.5), 1 volume of isopropanol, and 30 µg GlycoBlue (Invitrogen AM9516). Pellets were washed with 80% ethanol and resuspended in nuclease-free water. Ribosomal RNA was depleted using the siTOOLs Biotech riboPOOLs kit. Purified RPFs (1 - 5 µg) were hybridized to rRNA probes at 68°C for 10 min, then cooled to 23°C (0.1°C/sec). Hybridized samples were incubated with prepared streptavidin magnetic beads for 5 minutes at room temperature, then 5 minutes at 37°C. The rRNA-depleted supernatant was purified using the RNA Clean and Concentrator-5 kit (ZYMO RCC-5). Samples were denatured (95°C, 5 min) and resolved on a 15% TBE-urea polyacrylamide gel (Invitrogen EC68852) alongside a microRNA ladder and a custom 27/29/32 nt RiboCut marker. The gel was SYBR Gold-stained, and the 27-30 nt region was excised, disrupted, and extracted overnight at 4°C in 300 mM sodium acetate (pH 5.0), 0.05% SDS, and RNase inhibitor. Extracts were filtered (0.22 µm Costar Spin-X) and RNA was isopropanol-precipitated, washed, and resuspended.

(*3*) *<u>RPF library preparation:</u>* RPFs was end repaired with T4 PNK (NEB M0201) and 1 mM ATP (1 h, 37°C). RNA was ethanol-precipitated overnight with linear acrylamide, washed, and resuspended. Small RNA sequencing libraries were generated using the NEBNext Low-bias Small RNA Library Prep Kit (NEB E3420) according to the manufacturer’s protocol.

(*4*) *<u>RNA-seq library preparation:</u>* 10% combined lysate was extracted using TRIzol (Invitrogen 15596026) and the Direct-zol RNA Miniprep Plus Kit (ZYMO R2072) with on-column DNase I digestion. RNA-seq libraries were subsequently constructed from 50 ng of purified total RNA using the NEBNext Ultra II Directional RNA Library Prep Kit (NEB E7760), following the Poly(A) mRNA Magnetic Isolation Module (NEB E7490) for mRNA enrichment according to the manufacturer’s protocol.

### Global Translatome Data Analysis

*<u>(1) Ribo-seq and RNA-seq data processing:</u>* Raw Ribo-seq reads were first processed to remove 3’ adapter sequences (AGATCGGAAGAGCACACGTCTGAACTCCAGTCA) using Cutadapt, with reads shorter than 15 nucleotides subsequently discarded. To deplete ribosomal and non-coding RNA, the trimmed reads were first aligned to human ncRNA from Ensembl (GRCh38 release 115) (27) and the human 45S pre-rRNA sequence (NCBI GenBank U13369.1) using Bowtie (v1.1.2) (28) with parameters strictly constrained to a maximum of one mismatch and uniquely mapping reads only. Reads failing to map to this database were then aligned to a yeast spike-in reference (*Saccharomyces cerevisiae* cDNA, R64-1-1, Ensembl release 115) using identical stringent Bowtie parameters to quantify spike-in controls. Finally, the remaining unmapped reads representing human protein-coding transcripts were aligned to the human genome (GENCODE v32 primary assembly supplemented with tRNA annotations) (29) using STAR (v2.5.4a) (30). To ensure high-confidence footprint alignments, STAR was executed in end-to-end alignment mode with a maximum seed search start length of 15, restricting output to uniquely aligned reads with no more than two mismatches. For the associated RNA-seq data, raw paired-end sequencing reads were aligned directly to the human reference genome (GRCh38) using STAR, guided by the same GENCODE v32 annotation, generating coordinate-sorted alignments, transcriptome-coordinated alignments, and gene-level read counts. RNA-seq reads failing to map to the human genome were subsequently aligned to the yeast reference genome to quantify RNA-seq spike-in controls. All final coordinate-sorted BAM files across both pipelines were indexed using Samtools for downstream visualization.

*<u>(2) Ribo-seq QC and Differential Translation Efficiency (DTE) analysis:</u>* Ribo-seq quality control, empirical P-site identification, and extraction of coding sequence-mapped Ribosome Protected Fragments (RPFs) were performed using the riboWaltz R package (31). Specifically, P-site offsets were empirically calculated based on the metagene start codon periodicity for each specific read length. Transcript counts were subsequently aggregated to Ensembl Gene IDs via org.Hs.eg.db. For differential expression and translation efficiency (TE) analysis, gene-level RNA-seq and RPF counts were merged and analyzed using DESeq2 (32). To ensure robust statistical power and eliminate background noise from lowly expressed transcripts, an assay-specific expression filter was applied prior to normalization; only genes containing a minimum of 10 reads in at least two samples for both the RNA-seq and RPF datasets independently were retained. To control for global shifts in transcription and translation, manually calculated size factors were derived from the external yeast spike-in counts. As previously published (33), differential TE was evaluated using a multi-factor design modeling the statistical interaction between library assay (RNA vs. RPF) and treatment condition (TMZ vs. Control), defining significance as a Benjamini-Hochberg adjusted p-value < 0.05. Finally, genes exhibiting significantly downregulated translation efficiency (adjusted p-value < 0.05, log2 Fold Change < −1.0) were subjected to Gene Ontology over-representation analysis using the clusterProfiler R package (34) against a background universe of all detected genes.

*<u>(3) Correlation with sequence features:</u>* To assess the relationship between translation efficiency and various transcript features, differential TE metrics were integrated with corresponding RNA-seq expression data, sequence characteristics, and transcript stability profiles. RNA-seq baseline expression (baseMean) and expression log2 fold changes were extracted from DESeq2 outputs. Sequence-level features were derived from the human GRCh38 filtered CDS FASTA using the Biostrings R package. Specifically, overall CDS length, guanine (G) content percentage, and GC content percentage were computed for each transcript and averaged to yield gene-level consensus values. Canonical G-quadruplex (G4) motifs were identified within CDS regions via regular expression matching four tracts of at least three consecutive guanines, separated by loops of one to seven arbitrary nucleotides (35,36), and genes harboring one or more motifs across any transcript variant were categorized as G4- positive. Furthermore, mRNA decay rates were incorporated by mapping corresponding polysome entry MAP half-lives or cytoplasmic MAP half-lives from K562 TimeLapseSeq data (37). Statistical associations within significantly down-regulated TE targets (adjusted p-value < 0.05, log2 Fold Change ≤ −1.0) relative to background genes (defined as all remaining quantified genes failing to meet these combined significance and fold-change thresholds) were assessed via the ggpubr R package (38).

### Statistical Analysis

All image data was quantified using BioRad Image Lab software. Statistical analysis was performed using tests indicated in each figure legend. A 95% confidence level was set for every test with a definition of statistical significance at p < 0.05.

## Results

### Temozolomide-induced RNA alkylation impairs mRNA translation *in vitro*

To determine the direct impact of TMZ-induced alkylation damage on RNA function, we first established an *in vitro* system using synthetic firefly luciferase mRNA. The transcripts were exposed to 30 mM TMZ, vehicle control (DMSO), 30 mM MMS (positive control for RNA alkylation), or mock treatment for 30 minutes. Following treatment, the RNA was subjected to rigorous column purification for two rounds to remove residual chemical reagents (**Figure 1A**). Capillary electrophoresis confirmed that neither the incubation nor the chemical treatments induced significant transcript degradation, maintaining the expected 1,728 nucleotide full-length mRNA integrity (**Figure 1B**).

**Figure 1.**
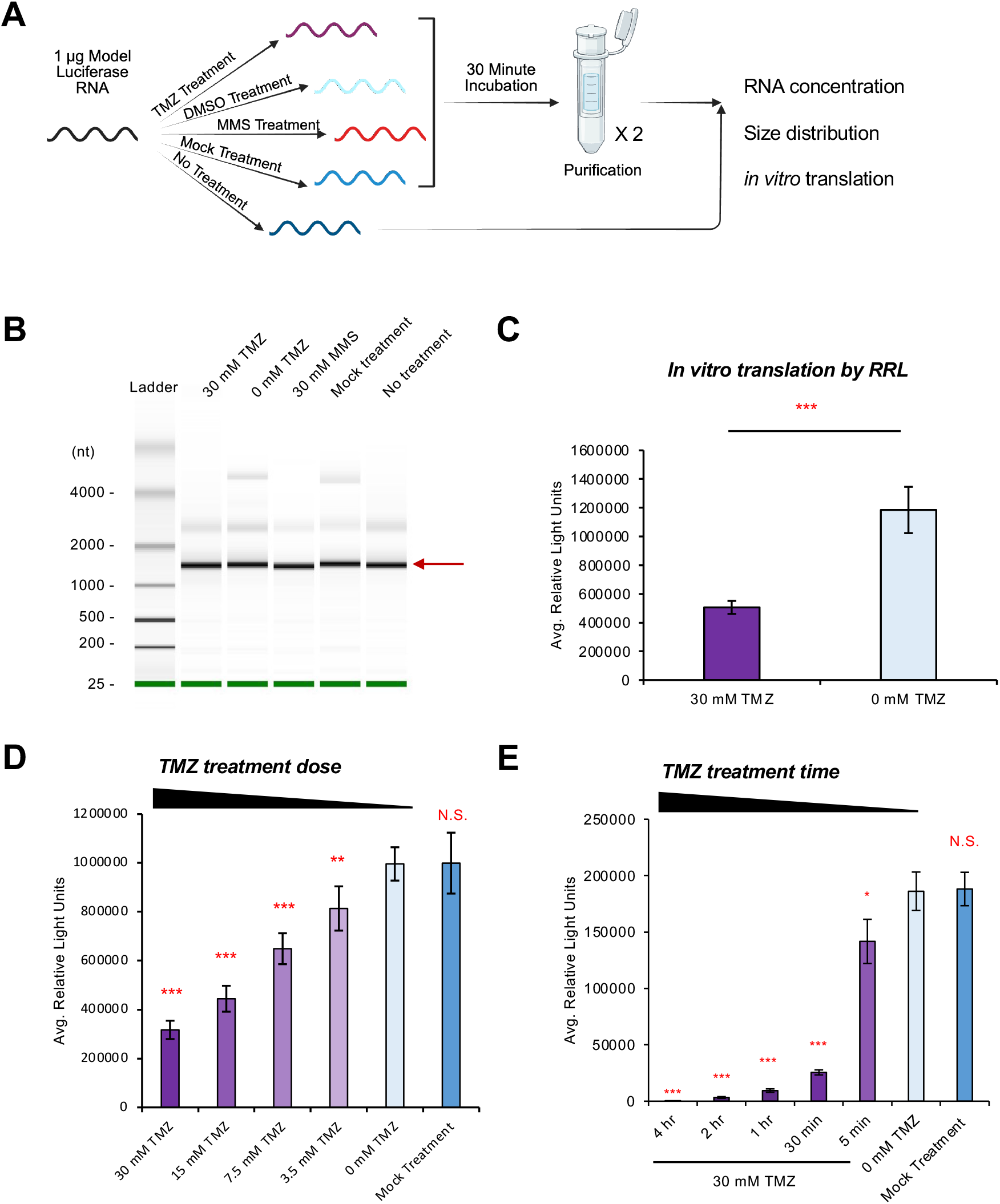
Temozolomide-induced RNA alkylation impairs mRNA translation *in vitro*. **(A)** Schematic of the *in vitro* treatment of firefly luciferase mRNA. RNA was exposed to TMZ, DMSO (vehicle), MMS (positive control) or mock treatment for 30 minutes, followed by two rounds of column purification. A no treatment control was included to account for potential RNA loss during incubation and purification. All samples were subsequently evaluated for concentration, size distribution and *in vitro* translation efficiency. **(B)** BioAnalyzer capillary electrophoresis confirming the integrity and size distribution of the treated RNA (full-length firefly luciferase mRNA = 1,728 nucleotides). **(C-E)** Translation efficiency of treated mRNAs assessed by rabbit reticulocyte *in vitro* translation assay. TMZ-treated mRNA displays significantly reduced translation activity compared to the DMSO control, demonstrating both (D) dose-dependent (0 – 30 mM) and (E) time-dependent (5 minutes to 4 hours) inhibition. Samples were treated with 30 mM for 30 minutes unless specified. Statistical significance was determined by two-tailed student’s t test (n = 3 independent replicates); * p < 0.05, ** p < 0.01, *** p < 0.001.

We next evaluated the functional consequences of this exposure using rabbit reticulocyte lysate *in vitro* translation assay. Transcripts treated with TMZ exhibited a profound and significant reduction in translation efficiency compared to DMSO-treated controls (**Figure 1C**). The degree of translational reduction for TMZ treatment was similar to MMS treatment (**Supplementary Figure 1**), which has been known to cause significant RNA alkylation (14–18). In addition, this translational inhibition was highly responsive to TMZ, demonstrating both a clear dose-dependent decline from 3.5 to 30 mM TMZ (**Figure 1D**) and a time-dependent decline from 5 minutes to 4 hours TMZ treatment (**Figure 1E**). Overall, these results support the hypothesis that TMZ treatment damages the mRNA translatability, most likely by direct alkylation.

### Temozolomide directly induces aberrant RNA methylation *in vitro*

Having observed severe translational impairment following TMZ exposure, we next sought to characterize the specific lesions responsible for this functional decline. To achieve robust absolute quantification via liquid chromatography-tandem mass spectrometry (LC-MS/MS), we focused our profiling on a targeted panel of alkylating modifications, specifically m^7^G, m^1^A, m^3^C, and m^6^A, for which validated analytical standards are available. m^6^G and m^3^A were not profiled due to the lack of available standards. As expected, control treatment has low basal modification levels (less than 0.1%). Following a 30-minute exposure to 30 mM TMZ, the *in vitro* treated luciferase mRNA exhibited a significant induction of specific alkylation lesions, most notably m^7^G (2%), alongside m^1^A (0.1%) and m^3^C (0.2%) modifications, when compared to the DMSO control (**Figure 2A-D**). This will be equivalent to an average of 7.9 m^7^G, 0.5 m^1^A and 0.7 m^3^C per molecule, based on the firefly luciferase RNA sequence. As a positive control that was known to induce RNA modifications, MMS strongly induced aberrant modifications, including m^7^G, m^1^A and m^3^C levels (**Supplementary Figure 2A-D**). For both TMZ and MMS, m^6^A level remained less than 0.001%, highlighting the specificity of such direct alkylation. To independently validate the mass spectrometry findings, we performed dot blot analyses, which confirmed a specific accumulation of m^7^G in both TMZ- and MMS-treated RNA substrates (**Figure 2E**). Together, these biochemical data demonstrate that TMZ directly deposits aberrant methyl groups onto single-stranded RNA, creating lesions that may physically interact with translational machinery.

**Figure 2.**
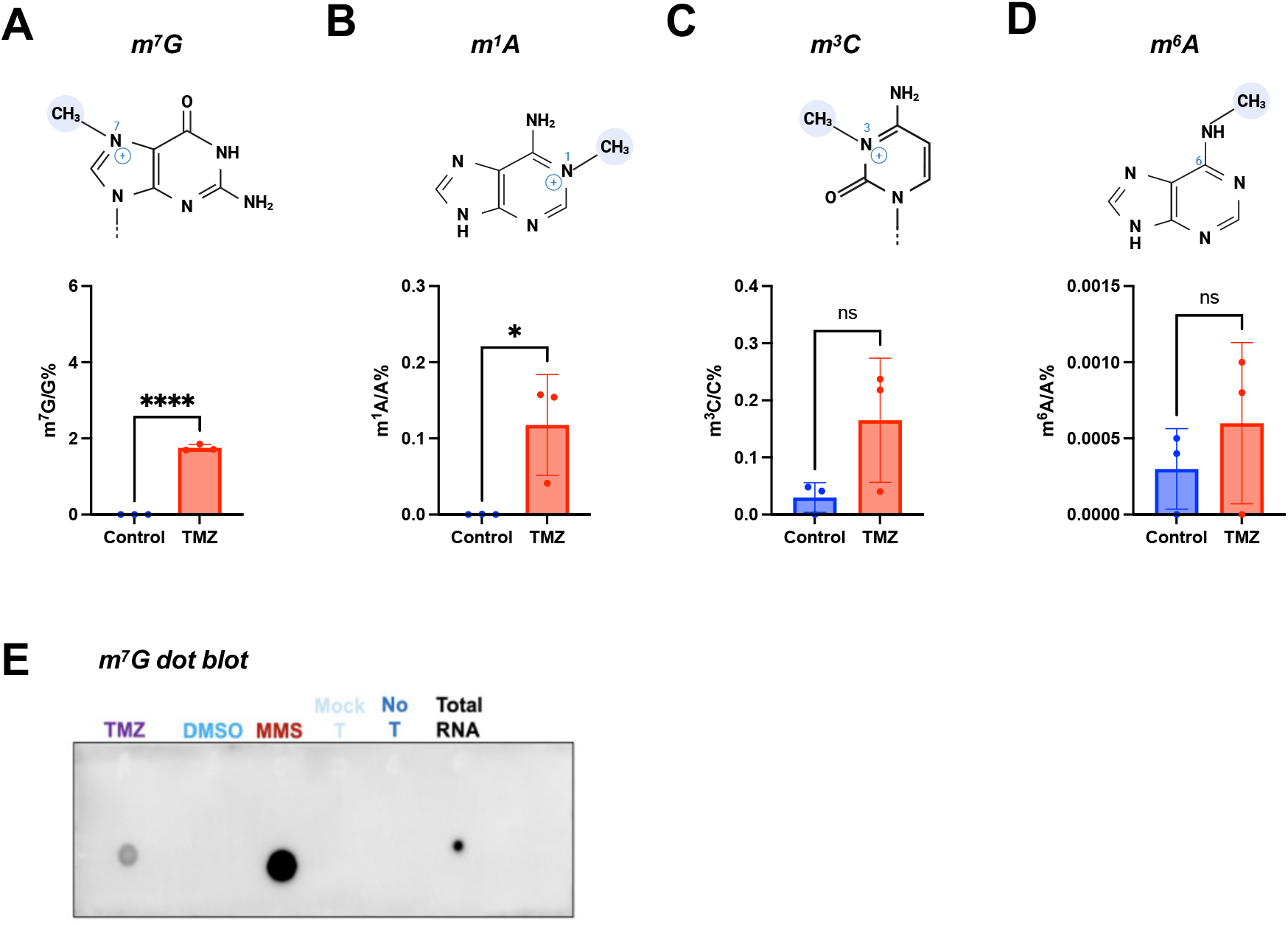
Temozolomide directly induces aberrant RNA methylation in vitro. **(A-D)** Mass spectrometry quantification of m^7^G, m^1^A, m^3^C and m^6^A levels in luciferase mRNA following a 30-minute *in vitro* treatment with 30 mM TMZ or DMSO control. For all MS data, modified nucleoside abundance is normalized to the respective unmodified nucleoside. **(E)** Dot blot analysis confirming the accumulation of m^7^G modifications following *in vitro* TMZ or MMS treatment. Statistical significance was determined by two-tailed student’s t test (n = 3 independent replicates); * p < 0.05, ** p < 0.01, *** p < 0.001.

### Acute temozolomide exposure induces RNA methylation and attenuates global protein synthesis in glioblastoma cells

To investigate whether the TMZ-induced RNA damage and subsequent translational repression observed *in vitro* (**Figure 1-2**) translates to a complex cellular environment, we evaluated global translational dynamics in U251-MG glioblastoma cells. To model the rapid pharmacokinetics of TMZ exposure *in vivo*, we employed an acute 1 mM treatment strategy. This allows for the robust biochemical capture of primary, direct RNA modifications independently of downstream apoptotic signaling. Total cellular RNA was extracted following a 2-hour acute exposure to 1 mM TMZ or 1 mM MMS. Consistent with our *in vitro* observations, mass spectrometry quantification of the cellular total RNA revealed a significant accumulation of aberrant modifications, primarily m^7^G, in response to both TMZ (**Figure 3A-D**) and MMS (**Supplementary Figure 3A-D**) treatments. Interestingly, in contrast to m^7^G, m^1^A and m^3^C did not show a TMZ-dependent induction; rather, we observed a modest but statistically significant decrease in these modifications following TMZ treatment. Given that m^1^A, m^3^C, and m^7^G are all highly enriched in native tRNAs, the specific increase in m^7^G strongly argues against a generalized upregulation of cellular tRNAs, pointing instead to direct chemical alkylation. Together, these data indicate that acute TMZ exposure in living cells produces aberrant RNA damage characterized at least in part by a significant accumulation of m^7^G.

**Figure 3.**
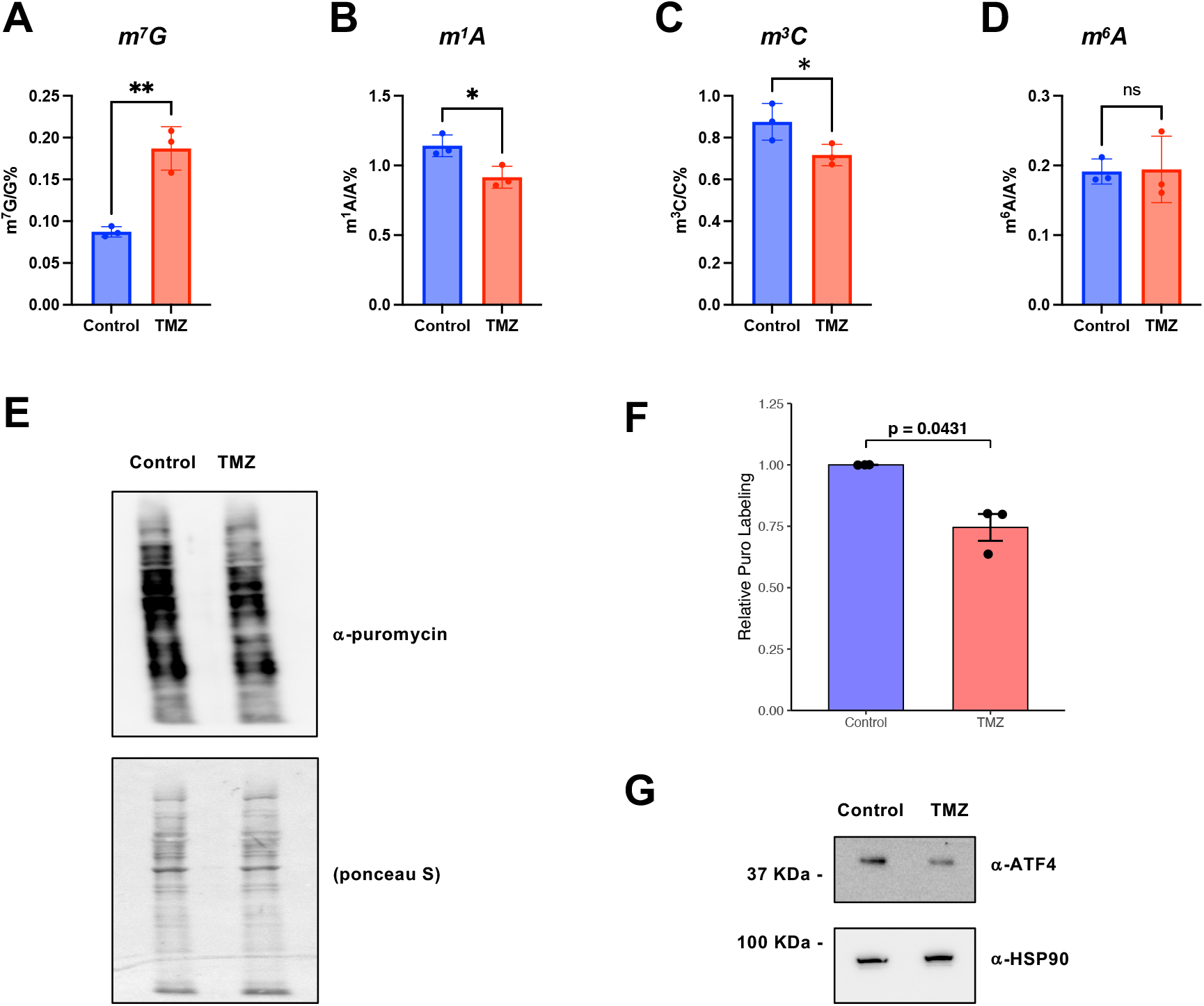
Acute temozolomide exposure induces RNA methylation and attenuates global protein synthesis in glioblastoma cells. **(A-D)** Mass spectrometry quantification of m^7^G, m^1^A, m^3^C and m^6^A in total cellular RNA from U251-GM cells. Cells were treated for 2 hours with 1 mM TMZ, alongside respective controls. Modified nucleosides were normalized to the unmodified counterparts. **(E-F)** Puromycin incorporation assay to detect global protein synthesis rates in U251-GM cells following 2 hours of 1 mM TMZ or DMSO Control, including **(F)** densitometric quantification of anti-puromycin western blots. **(G)** Western blot analysis of ATF-4 levels following acute TMZ treatment (2 hours, 1mM). HSP- 90 serves as the loading control. Statistical significance was determined by two-tailed student’s t test (n = 3 independent replicates); * p < 0.05, ** p < 0.01, *** p < 0.001.

We next assessed the immediate impact of this widespread RNA alkylation on cellular protein synthesis using a puromycin incorporation assay. Acute 2-hour treatment with TMZ or MMS triggered a statistically significant attenuation of global protein synthesis in the U251-MG cells (**Figure 3E-F**, **Supplementary Figure 3E-F**). To rule out the possibility that this translational repression was merely a secondary consequence of the Integrated Stress Response (ISR) triggered by generalized cellular toxicity, we evaluated the expression of ATF4, a classic downstream effector of the ISR. Western blot analysis revealed that ATF4 levels were not increased following 2-hour acute TMZ treatment (**Figure 3G**). This lack of ISR induction, together with the TMZ-reduced *in vitro* translation (**Figure 1**) strongly suggests that the observed reduction in global protein synthesis is likely a consequence of pervasive, TMZ-induced aberrant RNA methylation stalling the translational machinery.

### Nanopore direct RNA sequencing captures error signatures representing TMZ-mediated RNA damage

To evaluate aberrant RNA modifications at single-nucleotide resolution, we applied Nanopore direct RNA sequencing as an independent approach to overcome the limitations of LC-MS/MS. Because Nanopore sequencing measures ionic current variations caused by RNA modifications during translocation, modified residues frequently manifest as signature basecalling errors. We first sequenced *in vitro* TMZ- and Control-treated firefly luciferase mRNA. Guanine residues exhibited the highest frequency of basecalling errors upon TMZ treatment, which was predominantly driven by mismatches rather than insertions or deletions (**Supplementary Figure 4A–D**). To determine whether specific transcript locations are more susceptible to TMZ-mediated damage, we calculated the delta error by subtracting the control error rate from the TMZ-treated error rate. This analysis revealed 40 specific positions exhibiting a greater than 5% increase in error along the luciferase transcript (**Figure 4A**), which are dominated by guanines (32 positions). As a positive control to validate our experimental setup, we also sequenced MMS-treated luciferase RNA, which yielded a distinct damage landscape (**Supplementary Figure 4E**).

**Figure 4.**
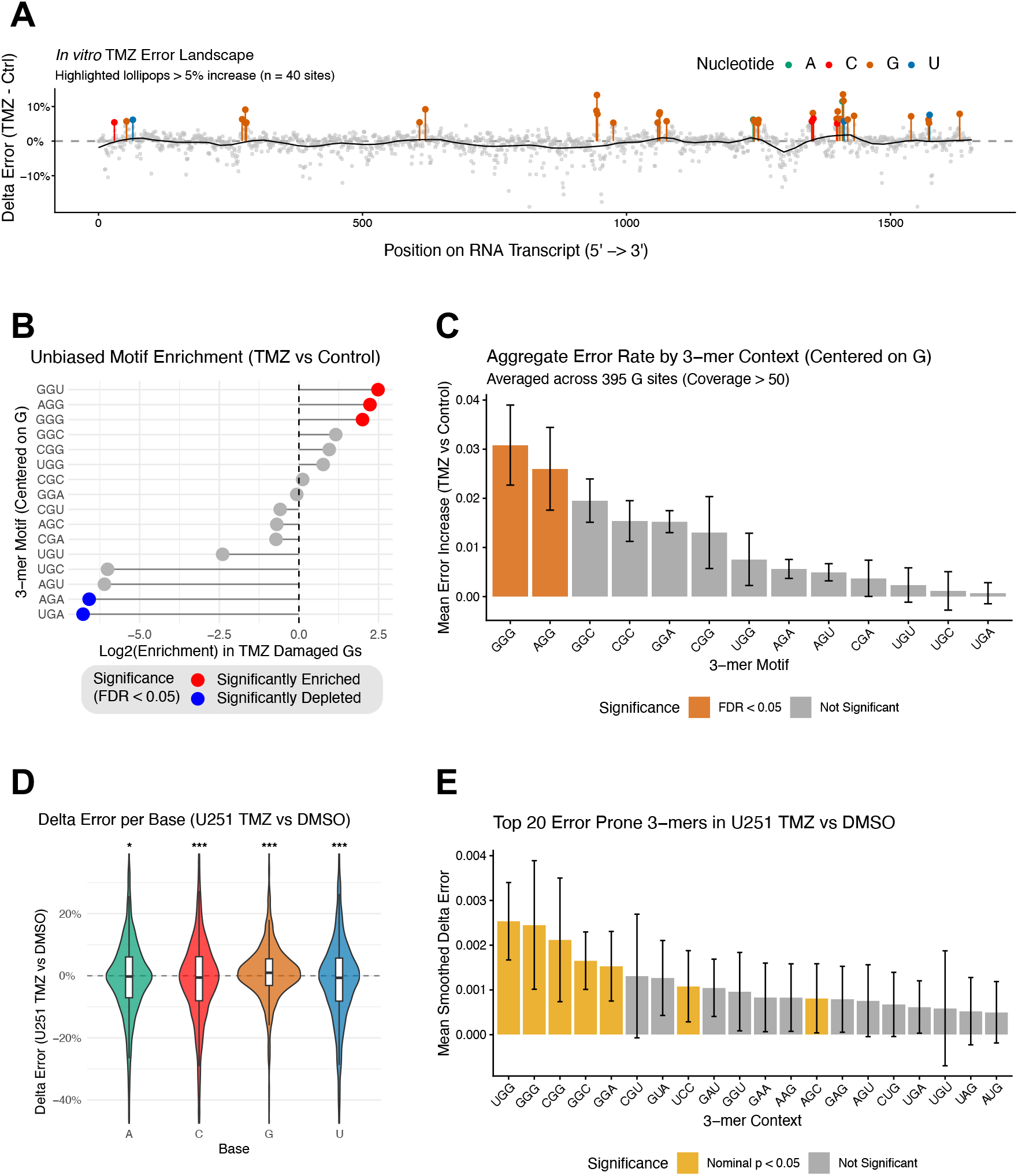
Nanopore direct RNA sequencing captures error signatures representing TMZ-mediated RNA damage. **(A)** Transcript-wide landscape plot of delta error from *in vitro* TMZ (30 mM, 30 minutes) versus control treatment across the firefly luciferase RNA sequence, with positions exhibiting a delta error greater than 5% highlighted. **(B)** Unbiased 3-mer motif analysis focused on Guanine (G) sites, comparing foreground sites with > 2% delta error and background sites with < 1% delta error, displayed as a lollipop chart where motif frequencies were compared using Fisher’s Exact Test with False Discovery Rate (FDR) correction. **(C)** Aggregate 3-mer analysis centered on Guanine positions with coverage greater than 50, where z-scores and FDR-adjusted p-values classify motifs into statistical tiers based on the mean mismatch increase between TMZ and control. **(D)** Violin and box plots showing delta error divided by base identity in TMZ-treated U251 cells (1 mM, 2 hours), evaluated via one-sample t-tests against a mean of zero (8,231 total sites from the top 10 expressed protein-coding transcripts, coverage cutoff > 50). **(E)** Analysis of U251 data using a 3-mer window to identify and display the top 20 3-mers exhibiting increased smoothed delta error based on z-scores and FDR adjustment.

Building on the recent observations that TMZ-induced genomic DNA damage displays distinct sequence motif preferences (11), we restricted our focus to guanine sites and performed an unbiased motif enrichment analysis. Comparing sites with an increased error rate greater than 3% against unmodified background sites with less than 1% change revealed GGU, AGG, and GGG as significantly enriched local sequence motifs (**Figure 4B**). To confirm these findings globally, we aggregated all 395 guanine sites and evaluated their surrounding 3-mer contexts, which similarly established that GGG and AGG motifs undergo significantly higher error rates upon TMZ exposure (**Figure 4C**).

Encouraged by these *in vitro* findings, we performed poly(A)-enriched Nanopore direct RNA sequencing on cellular RNA extracted from TMZ- versus DMSO-treated U251 glioblastoma cells. This generated approximately one million mapped reads, enabling a robust evaluation of highly expressed transcripts. Restricting our initial analysis to the top 10 expressed protein-coding transcripts, we evaluated error distributions across all four nucleotide bases and found that only guanine demonstrated a statistically significant increase in error rate following TMZ treatment (**Figure 4D**). Finally, expanding our scope to sequence context, we evaluated all local 3-mer windows across these transcripts to identify sequence-specific susceptibilities, revealing UGG, GGG, CGG, GGC, and GGA as the top five error-prone 3-mer sequences significantly induced by cellular TMZ exposure (**Figure 4E**). Together, these *in vitro* and cellular Nanopore direct RNA sequencing results establish a potential framework for detecting TMZ-induced RNA damage and uncovering underlying sequence-specific vulnerabilities.

### Quantitative translatome profiling reveals global translational repression and targeted vulnerability of proliferative networks by TMZ

Since endogenous internal m^7^G modifications have been previously shown to interact with specific reader proteins and perturb mRNA stability/localization, leading to lower gene expression (39,40), we hypothesized that the aberrant TMZ-induced m^7^G modification might directly interfere with translation. To this end, we sought to map this translational repression at transcript-level resolution. Because changes in Ribo-seq signal can result from either altered translation rates or underlying shifts in mRNA expression, we isolate true translational control by normalizing Ribo-seq against RNA-seq to determine translational efficiency (TE). Standard RNA-seq and Ribo-seq normalization methods often fail to capture global unidirectional shifts; therefore, we integrated a yeast lysate spike-in strategy to enable the accurate quantification of translational efficiency (TE) across the transcriptome. U251-MG cells were treated acutely with TMZ or a DMSO control, spiked with yeast lysate, and subjected to parallel RNA-seq and ribosome profiling (**Figure 5A**). Quality control analyses of the resulting ribosome-protected fragments (RPFs) confirmed the size enrichment of the Ribo-seq libraries, robust mapping to coding regions and expected metagene periodicity (**Supplementary Figure 5A-B**).

**Figure 5.**
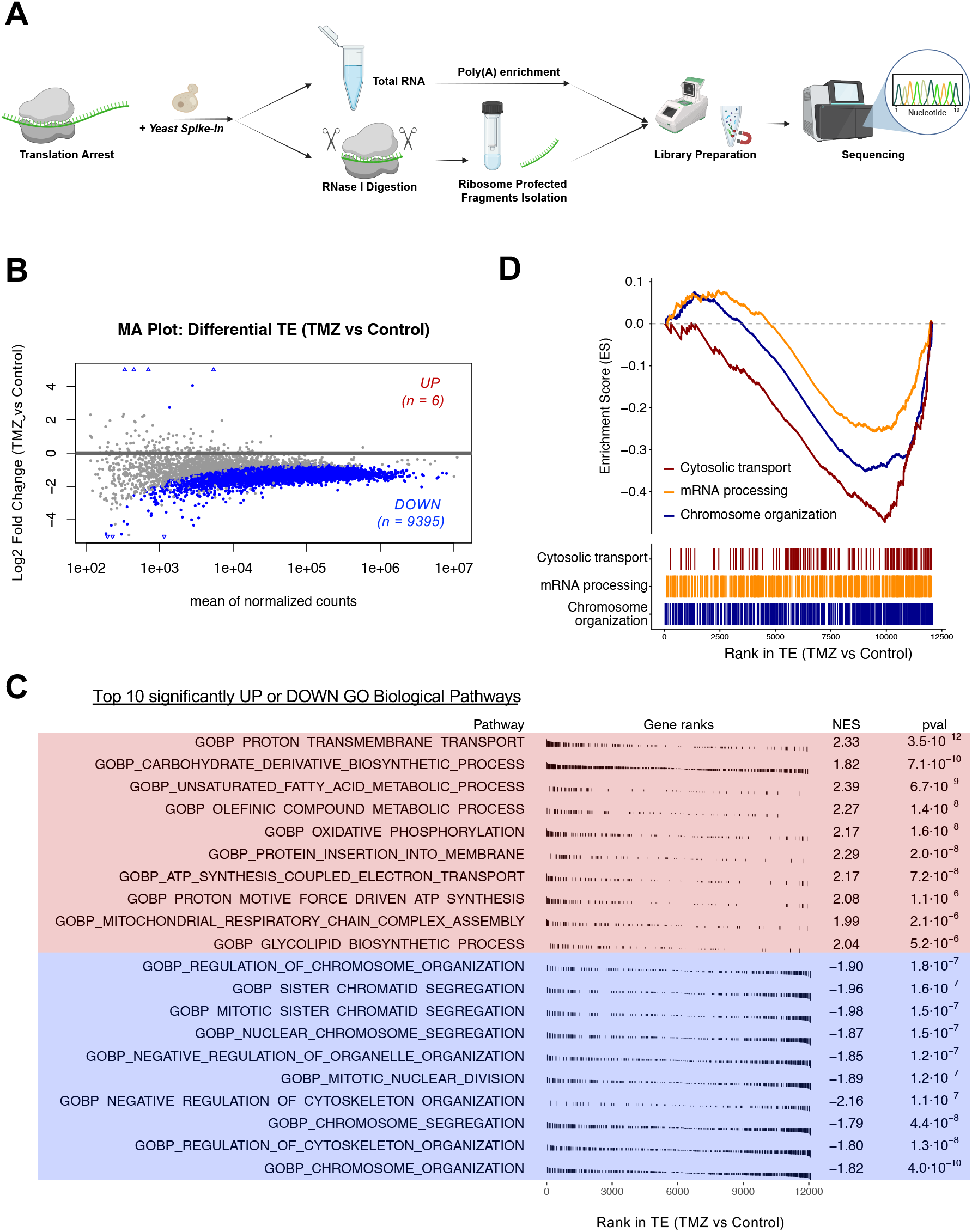
Quantitative translatome profiling reveals global translational repression and targeted vulnerability of proliferative networks by TMZ. **(A)** Experimental workflow for measuring global translational efficiency (TE) via Ribo-seq with a yeast spike-in. Cells were treated with 1 mM TMZ or a DMSO control for 2 hours and lysed in the presence of cycloheximide. A yeast lysate spike-in was added prior to RNase I digestion for normalization. *Top:* Total RNA was isolated and poly(A)-selected for RNA-seq library preparation. *Bottom:* Ribosome-protected fragments (RPFs) were purified for Ribo-seq library construction. **(B)** MA plot of yeast-normalized differential TE following acute TMZ treatment. The data demonstrates a global down-regulation of translation. Significantly altered transcripts (p.adj < 0.05) are highlighted in colored and counted. **(C)** Top 10 significantly up- or down-regulated GO Biological Pathways by Gene Set Enrichment Analysis (GSEA) of differential TE (TMZ vs Control), with genes exhibiting the highest differential TE fold change located at the far left. **(D)** Gene Set Enrichment Analysis (GSEA) of selected GO BP terms ranked by TE. The x-axis represents the complete dataset of genes ordered by decreasing TE by TMZ, with genes exhibiting the highest differential TE fold change (TMZ vs Control) located at the far left.

Yeast-normalized differential TE analysis revealed a massive, transcriptome-wide down- regulation of translation following acute TMZ treatment (**Figure 5B**), consistent with the overall decreased puromycin labeling (**Figure 3E-F**). Out of 11,411 detected genes, 9,395 showed a significant TMZ-induced reduction in TE and only 6 showed significant increase. The complete output of differential TE analysis can be found in **Supplementary Table 1**. TMZ treatment did not globally alter ribosome distribution across 5’ UTR, CDS and 3’ UTR regions (**Supplementary Figure 5A**), suggesting that the basic mechanics of translation initiation and termination remain largely intact, and pointing toward highly localized, sequence-specific translation elongation stalls.

We next investigated whether specific biological pathways were disproportionately affected by this widespread translational repression. Gene Set Enrichment Analysis (GSEA) of all transcripts ranked by their differential TE revealed a more severe translational repression on pathways related to chromosome organization and RNA processing, and less repression on pathways related to biosynthesis (**Figure 5C-D** and **Supplementary Table 2**). This selective vulnerability indicates that the TMZ-induced translational arrest disproportionately impairs critical cell cycle and chromosome organization networks, exerting an immediate, replication-independent stress on glioblastoma cells well before canonical DNA damage signaling occurs.

### Translational repression is driven by transcript guanine density and rapid initiation kinetics

To confirm that the observed TE down-regulation was driven by true translational repression rather than altered mRNA levels, we compared the log2 fold changes of mRNA abundance to their corresponding TE. This comparison demonstrated that the drop in translational efficiency occurred independently of both transcript abundance changes (**Figure 6A**) and basal RNA expression levels (**Supplementary Figure 6A**). This uncoupling suggests that TMZ is not targeting highly transcribed genes or altering mRNA stability.

**Figure 6.**
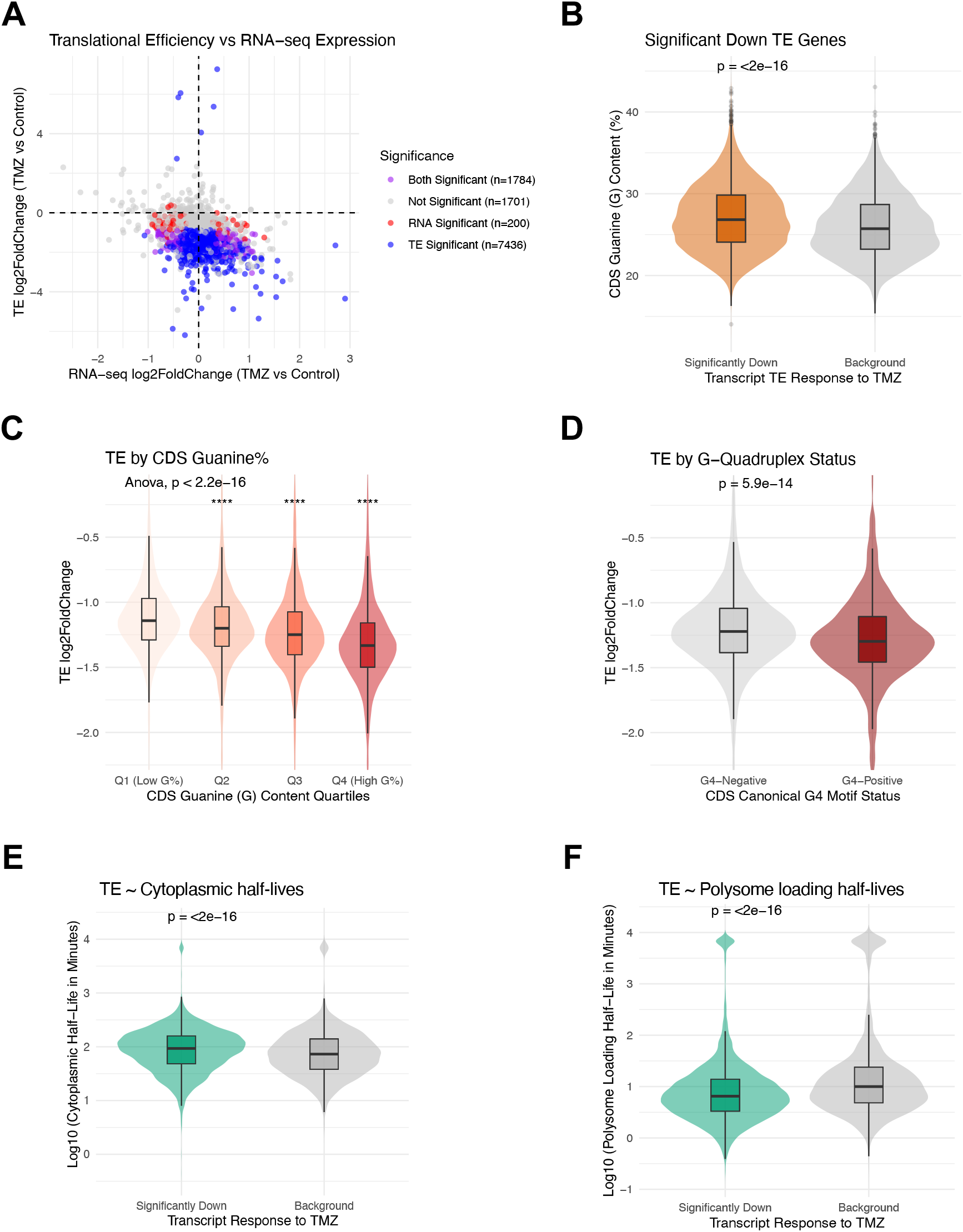
Translational repression is driven by transcript guanine density and rapid initiation kinetics. **(A)** Scatter plot comparing the log2 fold change of mRNA abundance (x axis) versus translational efficiency (y axis). Transcripts with statistically significant changes (p.adj < 0.05) are color-coded. **(B-C)** Down-regulated TE genes have higher guanine (G) content in the coding sequence (CDS). **(B)** Violin and box plot comparing the log2 fold change of translational efficiency (TE) following acute TMZ treatment, stratified by significantly down TE transcripts (p.adj < 0.05, Fold Change < −2) or background transcripts (remaining transcripts detected in Ribo-seq) **(C)** Violin and box plot displaying the log2 fold change of TE, stratified by CDS guanine content. Transcripts were binned into four equal quartiles (Q1-Q4) based on their absolute G-content percentage, with Q1 representing transcripts with the lowest guanine density and Q4 representing those with the highest. The central hinge represents the median, and the boxes span the first to third quartiles. Global statistical differences across all quartiles were assessed using an ANOVA test, and post- hoc comparisons were performed using two-sided Dunnett’s tests against Q1 group (p < 0.05, p < 0.01, *** p < 0.001, ns = not significant). **(D)** Violin and box plot comparing the translational efficiency (TE) log2 fold change of transcripts lacking canonical G4 motifs (G4-Negative) versus those containing at least one G4 motif within their coding sequence (G4-Positive). Statistical significance was determined using a two-sided Wilcoxon rank-sum test. **(E-F)** Violin and box plot comparing the **(E)** cytoplasmic half-live or **(F)** polysome loading half-life of transcripts significantly down-regulated following acute TMZ treatment (TE log2FC ≤ −1, adjusted p < 0.05) versus the unaffected background transcriptome. The y-axis displays the log10-transformed half-life in minutes derived from K562 TimeLapse-seq. Statistical significance was evaluated using a two-sided Wilcoxon rank-sum test.

We next sought to determine if the TMZ-induced lesions caused translational repression within specific sequence contexts. Given our biochemical findings that TMZ deposits m^7^G lesions onto single-stranded RNA, we hypothesized that the frequency of targetable guanines within the coding sequence (CDS) might dictate a transcript’s translational repression severity. Indeed, transcripts with significantly reduced TE exhibited significantly higher G-content (**Figure 6B**). Stratifying transcripts further demonstrated a clear dose-dependency, where transcripts with the highest guanine density exhibited the most severe down-regulation in translational efficiency compared to low-guanine transcripts (**Figure 6C**).

Since guanine-rich sequences are highly prone to forming G-quadruplex (G4) structures, which are known to act as stable hindrance to ribosome elongation (41,42), we also evaluated the presence of G4 motif in the CDS. As expected, transcripts containing canonical G4 motifs experienced significantly greater translational repression (**Figure 6D**). Importantly, however, this repression was driven primarily by absolute guanine density; elevated G-content reduced TE even in the absence of G4 motifs in the CDS (**Supplementary Figure 6B**). This suggests that the accumulation of individual alkylated guanines is more likely the driver of the translational repression. Furthermore, this effect was independent of overall CDS length (**Supplementary Figure 6C**) and was highly specific to the coding region, as neither guanine content nor G4 status in the 5’ UTR correlated with translational repression (**Supplementary Figure 6D-E**). Collectively, these data indicate that TMZ-deposited alkylation disproportionately stalls ribosomes on G-rich transcripts.

Having established that primary sequence composition dictates where physical TMZ lesions occur, we wondered if the dynamic lifecycle of these transcripts would affect the severity of the translational repression. Because chemical alkylation is a time-dependent process, we hypothesized that longer-lived transcripts would accumulate more damage and thus be more prone to translational repression. To this end, we evaluated baseline RNA decay and ribosome loading rates using subcellular TimeLapse-seq datasets (37). This method tracks metabolically labeled RNAs across fractionated cellular pools, enabling precise calculations of both transcript degradation kinetics and the rate at which newly synthesized RNAs are loaded onto ribosomes. Notably, a severe drop in translational efficiency correlated with a higher basal transcript stability in cytoplasmic compartment and whole cells (**Figure 6E** and **Supplementary Figure 6F**). This aligns with a cumulative exposure model, where increased transcript stability provides a longer temporal window for alkylation damage to accumulate.

We next evaluated the impact of translation initiation kinetics from subcellular TimeLapse-seq datasets (37). While rapid, dense ribosome loading could theoretically shield transcripts from alkylating agents, an assessment of polysome entry half-lives, a kinetic measure of translation initiation speed, demonstrated the opposite effect. Transcripts severely stalled by TMZ exhibited significantly shorter polysome entry half-lives compared to the background transcriptome (**Figure 6F**), indicating that they undergo rapid, high-frequency ribosome loading. Combined with our *in vitro* findings, these data suggest a synergistic mechanism of translational collapse. TMZ does not simply silence all stable RNAs; rather, the most profound repression occurs when stable, G-rich transcripts are simultaneously subjected to high rates of translation initiation. Because glioblastoma cells inherently rely on rapid, continuous polysome loading to sustain their growth, this rapid initiation actively drives ribosomes into accumulated alkylation roadblocks at high frequencies, culminating in a severe, targeted translational arrest.

## Discussion

In this study, we demonstrate that the chemotherapeutic agent temozolomide (TMZ) directly alkylates mRNA, leading to rapid translational repression. By integrating *in vitro* biochemical assays with quantitative yeast-spiked ribosome profiling, we uncoupled direct physical RNA damage from secondary cellular stress responses. Our data reveal that TMZ exposure induces significant accumulation of aberrant RNA modifications, which act as physical roadblocks to the translational machinery. This acute translational reprogramming is driven by transcript guanine density and rapid initiation kinetics, effectively repressing heavily translated, G-rich networks essential for chromosome organization and RNA processing.

The accumulation of aberrant methylations on mRNA fundamentally alters its biophysical properties and impairs interactions with the ribosome. While m^1^A, m^3^C, and m^7^G exist naturally often in tRNAs and rRNAs or as the defining feature of the 5’ mRNA cap, their stochastic accumulation within the coding sequence is highly deleterious. In mammalian cells, internal m^1^A, m^3^C, and m^7^G are kept at exceedingly low levels on mRNAs precisely because of their structure-disrupting nature. Alkylation at the *N*1 position of adenosine and the *N*3 position of cytosine directly disrupts the Watson-Crick base-pairing interface (21). This steric clash is well-documented to stall reverse transcriptases *in vitro* and is expected to obstruct the ribosomal decoding center if occurring on mRNA. On the other hand, alkylation at the *N*7 position of guanosine does not directly abrogate Watson-Crick base pairing. However, it introduces a positively charged moiety into the major groove of the RNA duplex, which can drastically alter local RNA folding dynamics and induce spontaneous depurination (20). Recent evidence demonstrates that internal m^7^G promotes mRNA degradation via IGF2BP proteins and triggers mRNA shuttling to stress granules by QKI proteins, effectively sequestering these transcripts away from the active translation pool and regulating cancer cell chemoresistance (39,40). Thus, TMZ-induced RNA alkylation likely paralyzes translation through both direct ribosomal collision and aberrant ribonucleoprotein (RNP) complex formation.

The spectrum of RNA damage we observed is intimately linked to the distinct chemical kinetics of the alkylating agents used. MMS operates primarily via an S_N_2 substitution mechanism, strongly favoring highly nucleophilic nitrogen atoms to generate m^7^G, m^1^A, and m^3^C lesions. In contrast, TMZ acts via an S_N_1 mechanism, decomposing spontaneously at physiological pH to form a highly reactive methyldiazonium cation. This intermediate is less sensitive to nucleophilicity, allowing it to rapidly alkylate both nitrogen and oxygen atoms. Crucially, our combined LC-MS/MS, Nanopore, and Ribo-seq cellular data establish guanines as the most vulnerable targets for this damage. This m^7^G accumulation aligns perfectly with recent independent antibody-based readouts, which similarly reported an increase in m^7^G and no change in m^6^A (43).

Nanopore direct RNA sequencing maps native epitranscriptomic modifications on intact transcripts (44). Previous algorithms and sequencing protocols have successfully identified endogenous RNA modifications like m^1^A, m^3^C, and m^7^G (45). Temozolomide deposits alkyation lesions onto nucleic acids. However, profiling these specific exogenous alkylation adducts directly on RNA remains poorly explored. Our data demonstrates that nanopore direct RNA sequencing can capture these aberrant RNA methylation events as basecalling errors (**Figure 4**). The sequencing output identifies distinct guanine-specific error signatures associated with TMZ damage. In DNA, temozolomide alkylation exhibits strong sequence context preferences (11). The alkylation process frequently targets guanine-rich motifs and specific non-canonical structural environments. Our analysis reveals analogous sequence context preferences for TMZ-induced damage in mRNA. This localized targeting dictates the physical distribution of RNA lesions. Consequently, this non-random damage distribution directly drives the transcript-specific translational repression observed in our ribosome profiling data.

While previous studies have shown that TMZ-induced DNA damage can eventually suppress protein synthesis through delayed secondary kinase signaling, such as AMPK-mediated mTORC1 inhibition or prolonged ISR (46,47), our data reveal a distinct immediate mechanism upon high dose TMZ treatment. Our findings align with an emerging paradigm that the efficacy and toxicity of classical DNA-targeting chemotherapeutics are significantly driven by their immediate impact on the transcriptome. For example, recent investigations into 5- azacytidine (5-AzaC), a cytidine analog traditionally characterized as a DNA methyltransferase inhibitor, have shown that the drug incorporates extensively into RNA co-transcriptionally (48). This 5-AzaC incorporation leads to aberrant RNA methylation patterns and subsequent disruptions in RNA stability and translation. Similar to our observations with TMZ, these non-canonical RNA-centric mechanisms challenge the dogma that alkylating and demethylating agents function solely through genomic DNA damage. Furthermore, recent studies highlight that rapid RNA alkylation serves as an early cellular alarm system; specifically, MMS-induced m^1^A deposition on RNA actively recruits DNA damage repair machinery to the sites of damage (14,18). Recognizing pervasive RNA alkylation highlights the urgent need to evaluate the under- appreciated mechanism of action for classic chemotherapies.

Although we observed a robust, dose-dependent correlation between CDS guanine density and translational repression, our footprint analysis did not reveal significant ribosomal pausing at specific G-rich sequence motifs. This apparent paradox is likely driven by the stochastic nature of TMZ-induced alkylation combined with cellular quality control mechanisms. Because TMZ modifies RNA stochastically, transcripts with higher G-content simply present a statistically larger target for random alkylation events. Consequently, ribosome stalling is distributed diffusely across many varying sites rather than concentrated at a single consensus sequence, creating a wide distribution of pauses that evades standard motif-finding algorithms.

In summary, we establish aberrant mRNA methylation as a targeted mechanism of translational collapse following TMZ treatment. However, several limitations must be considered. First, our experimental design utilized an acute high-dose TMZ exposure to robustly capture primary biochemical lesions and isolate direct translational repression before the onset of the ISR or widespread apoptosis. While this approach effectively uncoupled the mechanistic impact of RNA damage, future studies should evaluate the accumulation and translational consequences of these lesions under chronic, clinically relevant dosing regimens. Second, our mechanistic conclusions focus on the consequences of mRNA damage. Given that alkylating agents like MMS are well-documented to extensively modify tRNAs and disrupt their function (16), we cannot rule out the possibility that direct TMZ-induced alkylation of tRNAs or rRNAs acts in parallel to impair the translation machinery itself. Finally, overcoming the analytical limitations for other RNA alkylation not profiled here will be essential to understand the full RNA damage landscape. While our quantitative mass spectrometry robustly confirmed the accumulation of m^7^G, m^1^A, and m^3^C following *in vitro* TMZ exposure, the S_N_1 kinetics would predict that TMZ should also produce m^6^G and *N*3-methyladenine (m^3^A). Our study did not profile m^6^G and m^3^A directly due to the current lack of commercial standards required for accurate LC-MS/MS quantification and lack of commercial antibodies, representing a critical area for future investigation. Determining the precise coordinates of these lesions will clarify how local sequence context, beyond absolute guanine density, synergizes with translation initiation kinetics to drive therapeutic vulnerability.

## Data Availability

The Ribo-seq/RNA-seq generated in this study has been deposited in the Gene Expression Omnibus (GEO) database under accession code GSE338532. The Nanopore direct RNA sequencing data has been deposited in the Gene Expression Omnibus (GEO) database under accession code GSE343200.

## Supplementary Data statement

Supplementary Data include **Supplementary Figures 1-6** and **Supplementary Tables 1-2** in Excel format.

## Acknowledgements

We gratefully acknowledge the technical support and services provided by the Targeted Metabolomics and Proteomics Laboratory (TMPL) at the University of Alabama at Birmingham (UAB). Funds for the purchase of the SCIEX 7500+ Mass Spectrometer was provided by the University of Alabama at Birmingham Pathology Chair’s Office. We would like to thank Shane Rich-New and IT Research Computing Team at the University of Alabama at Birmingham for computational support. Schemes in Figure 1A, 5A and Graphical abstract were made with BioRender. We thank Yinsheng Wang (University of California at Riverside), Jun Zhang (University of Alabama at Birmingham) and Monima Anam (University of Alabama at Birmingham) for critical feedback on the manuscript. We thank Qi Chen (University of Utah) for helpful discussion.

## Author Contributions Statement

Conceptualization, J.R. and Z.S.; Resources, J.L.H. and Z.S.; Methodology, Investigation, Data Curation and Visualization, J.R., Z.S., A.J., K.M.B. and X.L.; Validation, A.J. and K.M.B.; Writing – Original Draft, Z.S. and J.R.; Writing – Review & Editing, J.R., A.J., X.L., K.M.B., J.L.H. and Z.S.; Funding Acquisition, Z.S., J.R. and K.M.B.; Supervision, Z.S.

## Supplementary Figure Legends

**Supplementary Figure 1.**
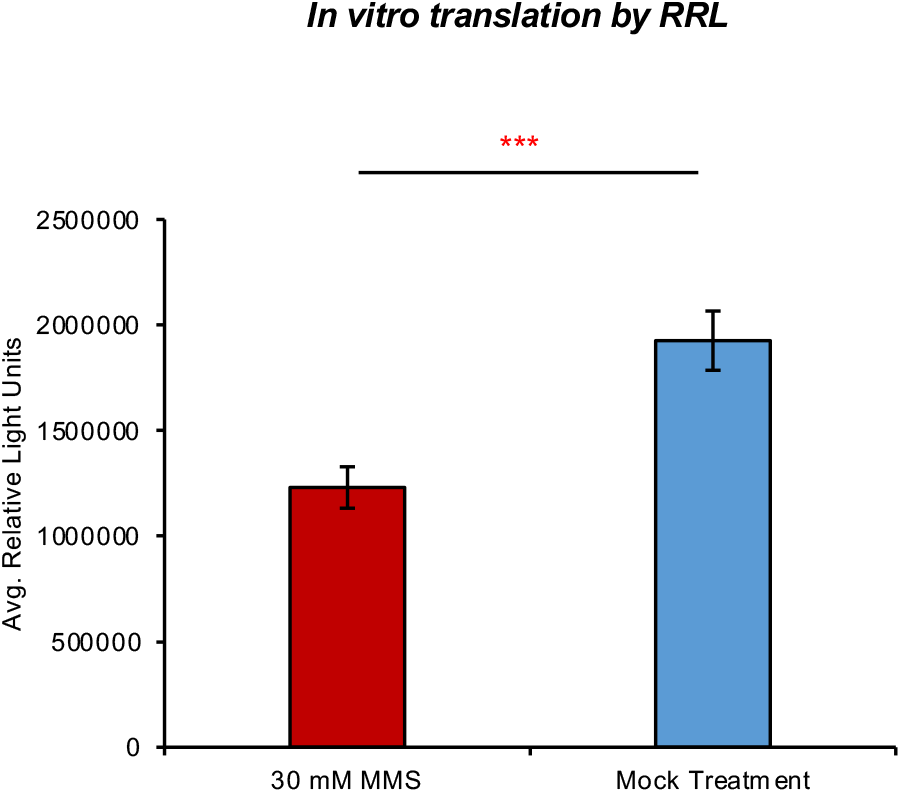
Temozolomide-induced RNA alkylation impairs mRNA translation *in vitro*. Translation efficiency of treated mRNAs assessed by rabbit reticulocyte *in vitro* translation assay. MMS-treated mRNA displays significantly reduced translation activity compared to the mock treatment control. Statistical significance was determined by two-tailed student’s t test (n = 3 independent replicates); * p < 0.05, ** p < 0.01, *** p < 0.001.

**Supplementary Figure 2.**
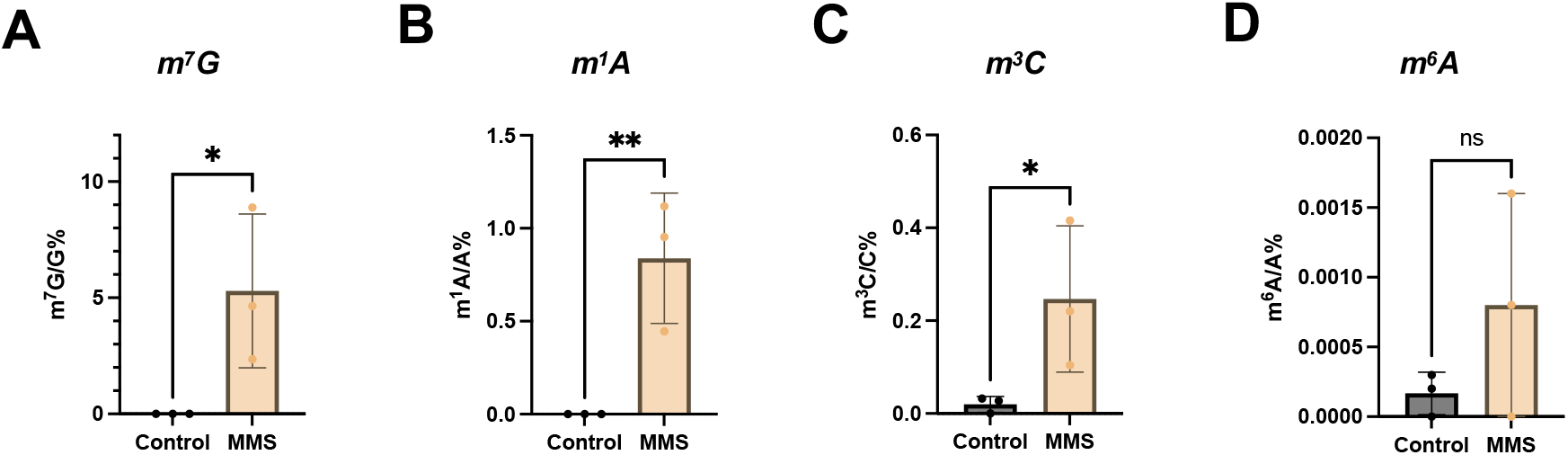
Temozolomide directly induces aberrant RNA methylation *in vitro*. **(A-D)** Corresponding mass spectrometry quantification of m^7^G, m^1^A, m^3^C and m^6^A in luciferase mRNA treated with 120 mM MMS or a mock treatment for 30 minutes *in vitro*. For all MS data, modified nucleoside abundance is normalized to the respective unmodified nucleoside. Statistical significance was determined by two-taxiled student’s t test (n = 3 independent replicates); * p < 0.05, ** p < 0.01, *** p < 0.001.

**Supplementary Figure 3.**
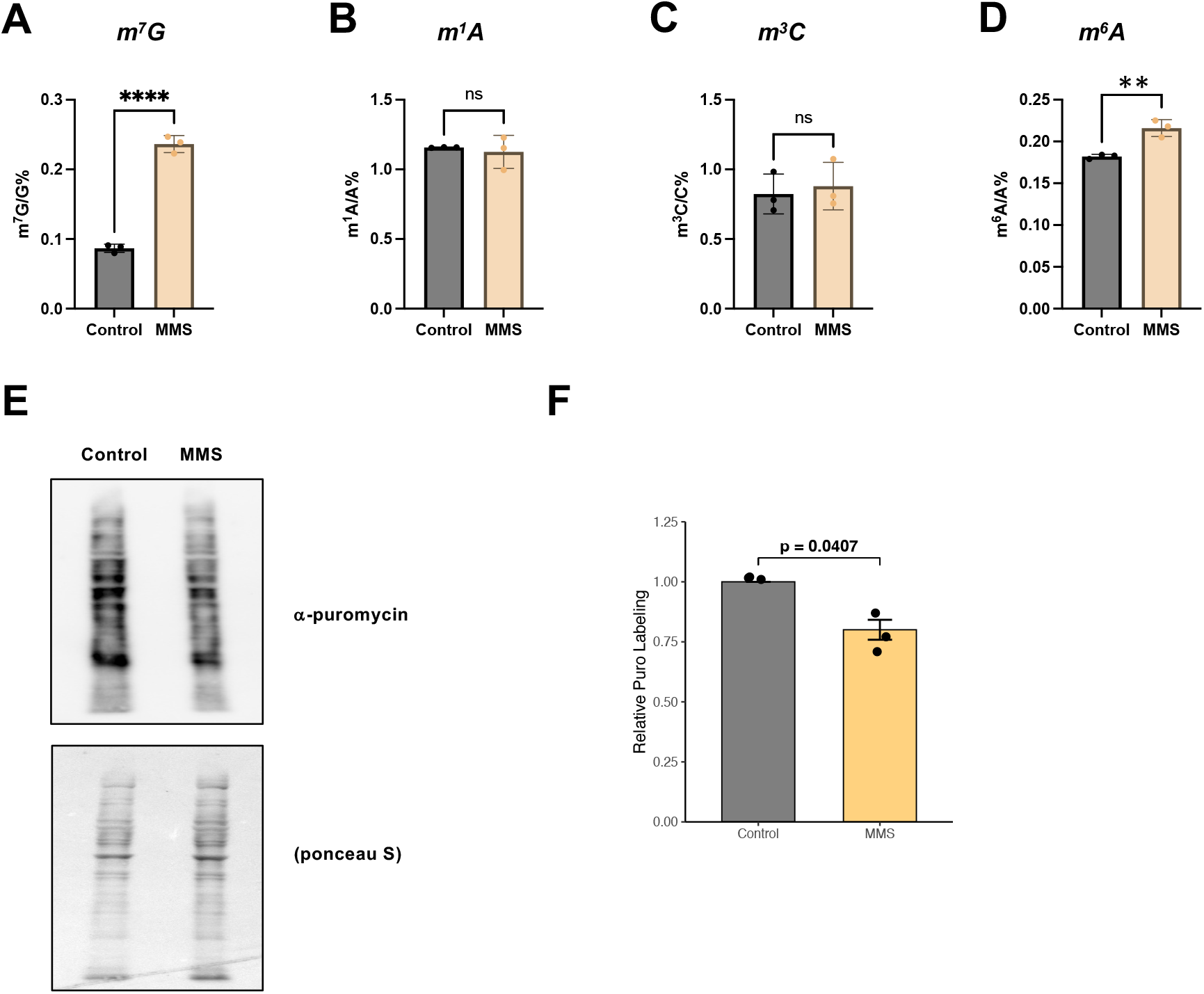
Acute temozolomide exposure induces RNA methylation and attenuates global protein synthesis in glioblastoma cells. **(A-D)** Mass spectrometry quantification of m^7^G, m^1^A, m^3^C and m^6^A in total cellular RNA from U251-GM cells. Cells were treated for 2 hours with 1 mM MMS, alongside respective controls. Modified nucleosides were normalized to the unmodified counterparts. **(E-F)** Puromycin incorporation assay to detect global protein synthesis rates in U251-GM cells following 2 hours of 1 mM MMS or Mock treatment Control, including **(F)** densitometric quantification of anti-puromycin western blots. Statistical significance was determined by two-tailed student’s t test (n = 3 independent replicates); * p < 0.05, ** p < 0.01, *** p < 0.001.

**Supplementary Figure 4.**
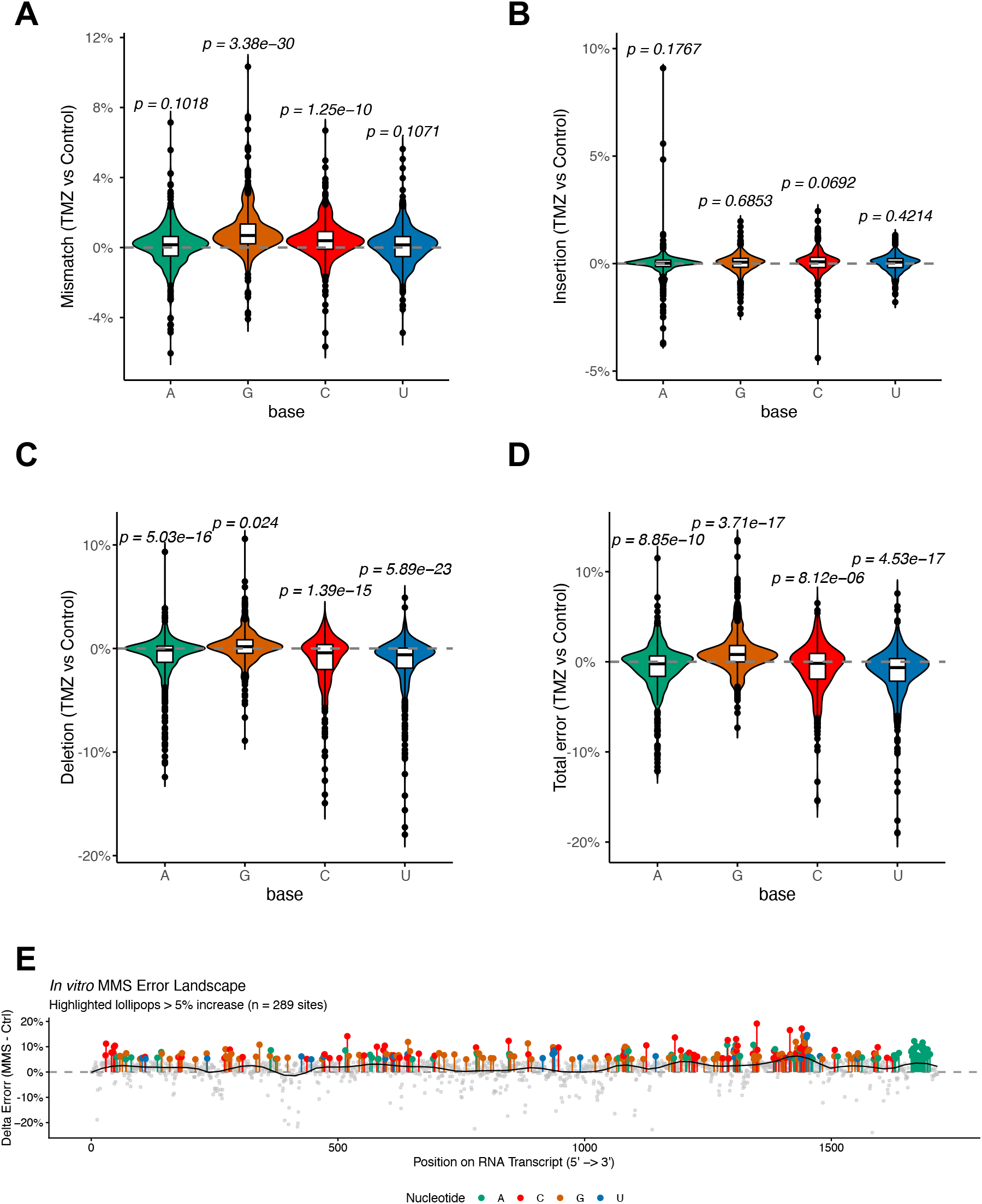
Nanopore direct RNA sequencing captures error signatures representing TMZ-mediated RNA damage. **(A–D)** Violin and box plots showing delta error grouped by base identity for (A) mismatch, (B) insertion, (C) deletion, and (D) total error based on *in vitro* TMZ versus control treatment datasets, evaluated via one-sample t-tests against a mean of zero. **(E)** Transcript-wide landscape plot of delta error from *in vitro* MMS (120 mM, 30 minutes) versus control treatment across the firefly luciferase RNA sequence, with positions exhibiting a delta error greater than 5% highlighted.

**Supplementary Figure 5.**
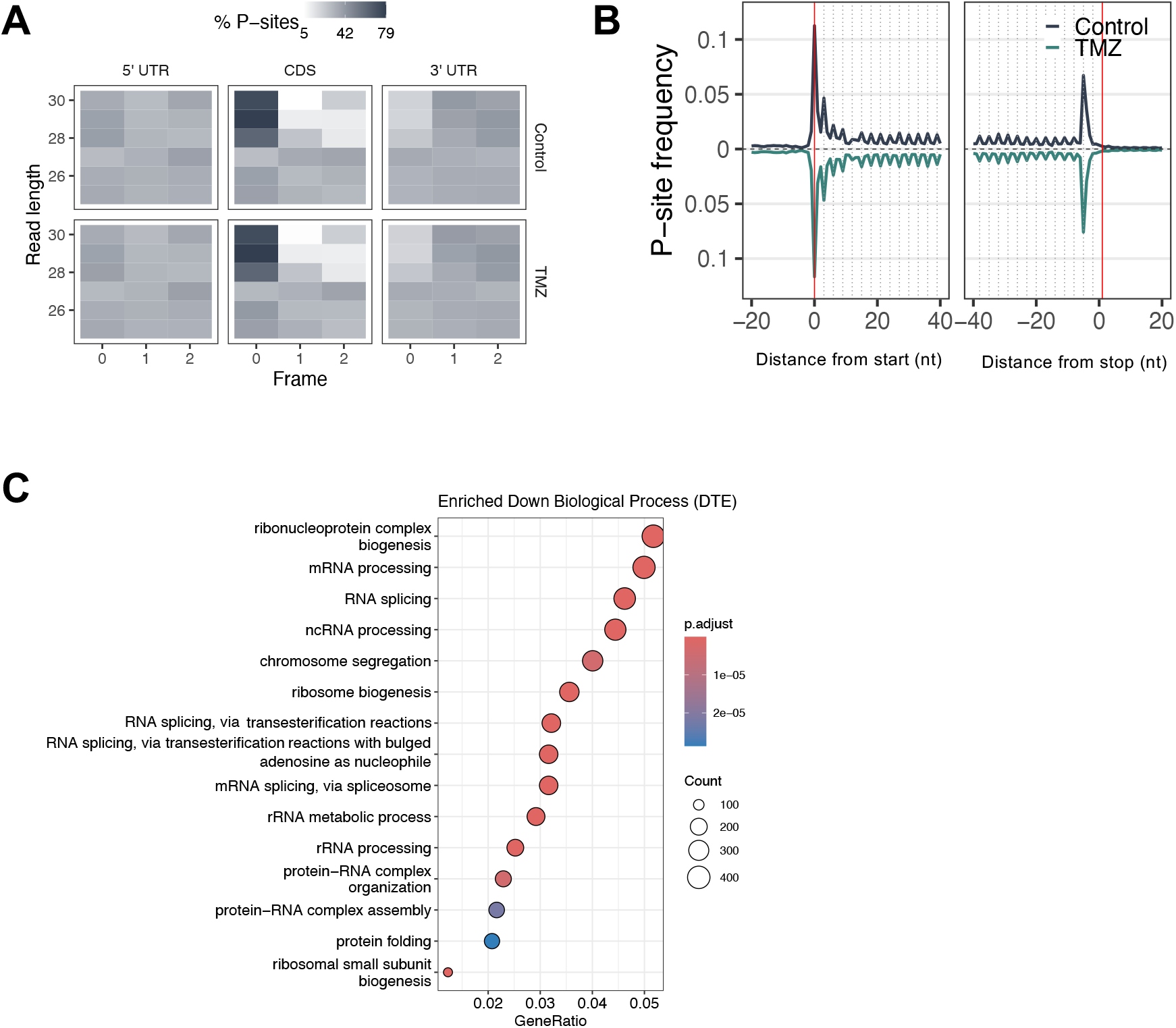
Quantitative translatome profiling reveals global translational repression and targeted vulnerability of proliferative networks by TMZ. **(A-B)** Ribo-seq quality check displaying the distribution of mapped P-sites across distinct transcript regions (A) and the metagene profile of P-site density relative to the start and stop codons (B). **(C)** Gene Ontology (GO) over-representation analysis of transcripts with significantly down- regulated TE (p.adj < 0.05, Fold Change < −2). The top 15 enriched Biological Pathway (BP) terms are displayed.

**Supplementary Figure 6.**
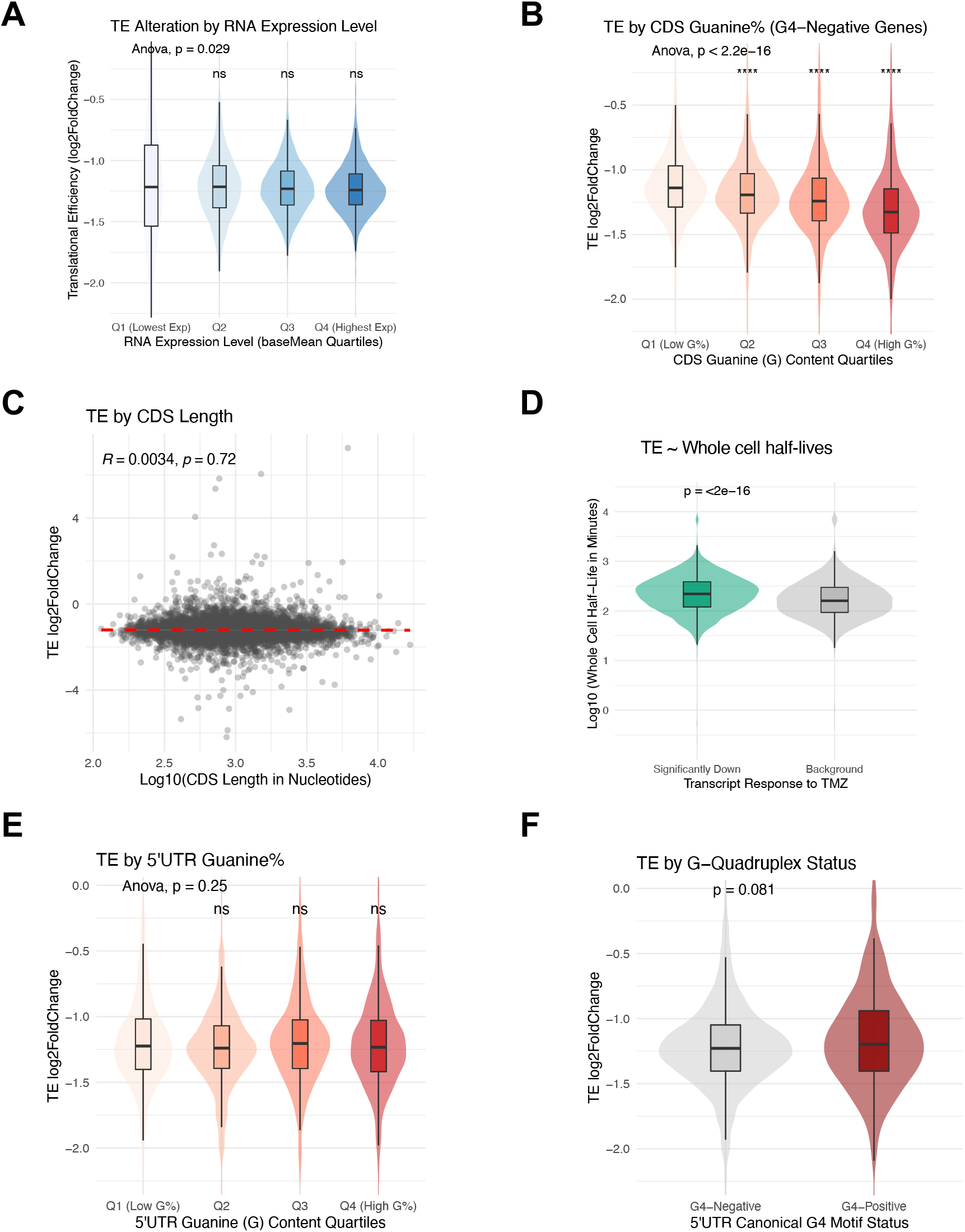
Translational repression is driven by transcript guanine density and rapid initiation kinetics. **(A)** Violin and box plot displaying the log2 fold change of translational efficiency (TE) following acute TMZ treatment, stratified by basal transcript abundance. Transcripts were binned into four equal quartiles (Q1-Q4) based on their mean normalized RNA-seq counts (baseMean), with Q1 representing the lowest expression and Q4 the highest. The central hinge represents the median, and the boxes span the first to third quartiles. **(B)** Down-regulated TE genes have higher guanine (G) content in the coding sequence (CDS) excluding G4-containing genes. Violin and box plot displaying the log2 fold change of TE, stratified by CDS guanine content. Transcripts were binned into four equal quartiles (Q1-Q4) based on their absolute G-content percentage, with Q1 representing transcripts with the lowest guanine density and Q4 representing those with the highest. Global statistical differences across all quartiles were assessed using an ANOVA test, and post-hoc comparisons were performed using two-sided Dunnett’s tests against Q1 group (p < 0.05, p < 0.01, *** p < 0.001, ns = not significant). **(C)** TMZ-induced translational repression does not correlate with transcript length. Scatter plot assessing the relationship between coding sequence (CDS) length and translational efficiency (TE) alteration. The x-axis displays the log10-transformed CDS length in nucleotides, and the y-axis represents the TE log2 fold change following acute TMZ treatment. The red dashed line indicates the linear regression fit. Statistical correlation was determined using a Spearman correlation test (R and p-values are displayed). **(D)** Violin and box plot comparing the whole cell half-life of transcripts significantly down-regulated following acute TMZ treatment (TE log2FC ≤ −1, adjusted p < 0.05) versus the unaffected background transcriptome. The y-axis displays the log10-transformed half-life in minutes derived from K562 TimeLapse-seq. Statistical significance was evaluated using a two-sided Wilcoxon rank-sum test. **(E-F)** Down-regulated TE genes do not have higher guanine (G) content in the 5’ UTR sequence. **(E)** Violin and Box plot displaying the log2 fold change of TE, stratified by 5’ UTR guanine content. Transcripts were binned into four equal quartiles (Q1-Q4) based on their absolute G-content percentage, with Q1 representing transcripts with the lowest guanine density and Q4 representing those with the highest. Global statistical differences across all quartiles were assessed using an ANOVA test, and post-hoc comparisons were performed using two-sided Dunnett’s tests against Q1 group. **(F)** Violin and box plot comparing the translational efficiency (TE) log2 fold change of transcripts lacking canonical G4 motifs (G4-Negative) versus those containing at least one G4 motif within their 5’ UTR sequence (G4-Positive). Statistical significance was determined using a two-sided Wilcoxon rank-sum test.

